# Impact of water deficit on single grapevine berry ripening

**DOI:** 10.1101/2023.09.18.557818

**Authors:** Mengyao Shi, Stefania Savoi, Thierry Simonneau, Agnès Doligez, Florent Pantin, Laurent Torregrosa, Charles Romieu

## Abstract

The effect of water deficit on grapevine fruit ripening has most often been addressed under the assumption that individual berries behave identically to their blend in the future harvest, both kinetically and metabolically. However, mixing unsynchronized berries, whose water and sucrose import pathways critically change according to their own developmental stages, intrinsically blurs the physiological and phenological effects of stress. We investigated the consequences of water deprivation on berry growth and primary metabolites content (glucose, fructose, tartaric and malic acids) on sixteen genetically distant genotypes of *Vitis vinifera* and fungus-tolerant hybrids submitted to 10 watering regimes, from well-watered to partial leaf shedding. Then, six genotypes were selected for comprehensive single berry analyses. Own-rooted potted plants bearing berries at the late herbaceous plateau stage were subjected to the different water treatments for four weeks in a greenhouse with automated regulation of soil water content. Berry and cluster growth were monitored by image analysis, before performing a final destructive sampling to determine berry weight and composition. Grape phenology was highly dependent on water availability. In some cultivars, ripening was considerably delayed or even prevented under well-watered conditions. These cultivars required an intermediate water deficit to trigger the second berry growth period along with sugar accumulation and malate breakdown, typical of the ripening process. Ripening still occurred in all genotypes upon severe water deprivation, although sugar accumulation and concentration were dramatically impaired, and the second growth period was annihilated or even replaced by shrivelling. Water deficit increased malate breakdown, uncoupling it from sugar accumulation. Single berry analyses suggest that although asynchronicity in berry ripening was reduced upon stress, individual fruits within the same cluster may undergo heterogeneous water budgets, expansion or shrivelling.

## Introduction

The response of grapevine (*Vitis vinifera* L.) vegetative development, productive performance, metabolism, and berry composition to water deficit (WD) has been extensively studied on many combinations of rootstocks, cultivars, and climate conditions (Lavoie-Lamoureux *et al*., 2017; López *et al*., 2007; Suter *et al*., 2019; reviewed in Gambetta *et al*., 2020). Studies over the past two decades have provided important information on WD effects on berry weight, berry total soluble solids (TSS) or sugars, titratable acidity (TA), pH, organic acids, phenolics, anthocyanins and terpenoids (des Gachons *et al*., 2005; Bindon *et al*., 2008; Leeuwen *et al*., 2009; Savoi *et al*., 2016, 2017). A number of experiments suggest that many factors, including soil, climate, vineyard management, and genotype, can influence vine response to WD, as reviewed by Lavoie-Lamoureux *et al*. (2017), Osakabe *et al*. (2014) and Ollat *et al*. (2019). The effects of WD largely depend on its timing and intensity during vine development. Early WD modifies berry set, phenological stages, and berry growth (Castellarin *et al*., 2007). WD at pre-véraison stages induces major modifications in malate, tartrate, glucose, and fructose concentrations. Berry volume remains small even after re-watering and is irreversibly affected by early WD (Keller *et al*., 2015; Keller & Shrestha, 2014; Korkutal *et al*., 2011; Shellie, 2014). Comparing eight irrigation scenarios, (Mccarthy, 1997) observed variable decrease in maximum berry weight upon stopping irrigation, depending on the timing of water stress and on the vintage. Ojeda *et al*. (2015) imposed three WD levels on cv. ‘Syrah’ and observed a subsequent reduction in berry growth, whose magnitude critically relied on the timing of stress application and its severity. Noticeably, shrunken pre-véraison berries suddenly regained their turgidity and expanded again before rewatering, when starting to ripen (Keller *et al*., 2015), due to the decrease of vacuolar osmotic pressure (Matthews *et al*., 1987) provoked by the induction of the apoplasmic pathway of phloem unloading (Savoi *et al*., 2021; Zhang *et al*., 2006). After véraison, WD effects were more variable, increasing the time necessary to reach the maximum berry volume that no longer doubled during ripening (Ojeda *et al*., 2015). However, berry volume can recover after re-watering (Girona *et al*., 2009; Intrigliolo & Castel, 2008; Munitz *et al*., 2017).

Generally, a medium WD after véraison, characterised by a predawn leaf water potential (Ψb) ranging from −0.4 to −0.6 MPa, reduces final berry weight and titratable acidity but increases TSS, phenolics, and total anthocyanins concentration, which can be associated with improved phenolic quality (Castellarin *et al*., 2007; Geng *et al*., 2022). In water-limited areas, the deficit irrigation method is used to manipulate berry composition and improve wine sensory properties (Chapman *et al*., 2005; Savoi *et al*., 2020) Nevertheless, these beneficial effects disappear when WD becomes severe (Ψb < −0.6 MPa) (Farooq *et al*., 2009; Seleiman *et al*., 2021). The increase in TSS is frequently interpreted as an acceleration of sugar accumulation in berries (Castellarin *et al*., 2007; Geng *et al*., 2022). However, WD impact on the respective fluxes of water and solutes accumulation by the fruit has barely been addressed (Liu *et al*., 2007) and there is frequent confusion between sugar accumulation in berries, which depends on sugar loading and metabolism, and sugar concentration, which also depends on water fluxes, berry growth and the resulting dilution. In addition, the phenological and metabolic responses of the fruit are inevitably confused in usual, averaged samples of non-synchronized berries (Shahood *et al*., 2020). This confusion should be particularly serious regarding water stress effects on water and sugar fluxes since xylem and phloem fluxes display opposite variations according to the fruit phenological stage (Ollat *et al*., 2019), before stopping in over-ripening, shrivelling berries (Savoi *et al*., 2021). Consequently, despite the huge amount of work on the effects of water status on grape yield and composition, no clear relationship could be established at the berry level between water flux, berry growth, sugar accumulation and concentration (Mirás-Avalos & Intrigliolo, 2017). The aim of the present work is to address WD effects on berry growth and changes in composition during ripening, not only considering the average of population but also single berries.

## Materials and methods

### 1. Plant material and growth conditions

Sixteen genotypes of *Vitis vinifera* and new disease-tolerant interspecific hybrids were investigated, containing 9 red grape genotypes (‘23298Mtp7’, ‘Affenthaler’, ‘7G20’, ‘VDQAG14’, ‘7G31’, ‘7G39’, ‘Mission’, ‘Staphidampelo’, ‘Vinhão’) and 7 white grape genotypes (‘Arbane’, ‘Savagnin blanc’, ‘Opsimo Edessis’, ‘VDQAG5’, ‘23298Mtp40’, ‘Cococciola’, ‘Garrido Macho’). These genotypes included 9 cultivars selected from a larger diversity panel (Nicolas *et al*., 2016) to represent the largest possible diversity (‘Arbane’, ‘Staphidampelo’, ‘Savagnin blanc’, ‘Vinhão’, ‘Affenthaler’, ‘Garrido Macho’, ‘Opsimo Edessis’, ‘Mission’, ‘Cococciola’) (https://bioweb.supagro.inra.fr/collections_vigne/Home.php), 5 offsprings from the ‘Syrah’ x ‘Grenache’ F1 progeny (‘23298Mtp7’, ‘2398Mtp40’, ‘7G20’, ‘7G31’, ‘7G39’) (Adam-Blondon *et al*., 2004), and 2 *Muscadinia rotundifolia* x *V*. *vinifera* disease-tolerant hybdrids known to display a reduced sugar content at ripe stage (‘VDQAG5’, ‘VDQAG14’) (Bigard *et al*., 2022). Potted, own-rooted two-year-old plants were initially grown under full irrigation from bud break to a few days before the date of véraison, as expected according to previous phenological records. Plants were then transferred to Phenodyn (https://www6.montpellier.inrae.fr/lepse_eng/Phenotyping-platforms/ Montpellier-Plant-Phenotyping-Platforms-M3P/PhenoDyn), a high-throughput phenotyping platform where air temperature and humidity are permanently regulated. Upon entrance, plants were subjected to a continuum of 10 watering regimes (one plant per regime and per genotype; Table 1) ranging from well-watered (treatment a, 2.4 g water g^−1^ dry soil) to severe water deficit generating partial leaf shedding (treatment j, 0.4 g water g^−1^ dry soil). Potted plants were continuously weighed and automatically watered every day to keep soil water content close to the target in each pot. The soil substrate was KL BP2 Base 274 + Argile 70L, including 170 kg m^−3^ clay, peat (50% frozen black peat, 50% white peat 0-20 mm) (Klasmann Deilmann, France). Average day and night temperatures and VPD were 30.4 °C, 22.8 °C, 1.96 and 1.01 kPa respectively, with a maximum PAR of 1280 µmol/m2/s.

**Table 1:** Codes used for the water treatments.

| Treatment | a | b | c | d | e | f | g | h | i | j |
| --- | --- | --- | --- | --- | --- | --- | --- | --- | --- | --- |
| soil water content (g water g<sup>-1</sup> dry soil) | 2.4 | 2.1 | 1.8 | 1.5 | 1.2 | 0.9 | 0.7 | 0.6 | 0.5 | 0.4 |

#### 1.2. Sample and data collection

Clusters were harvested on a first set of five plants per genotype, dedicated to the onset of WD (T0), a few days before the véraison date, as anticipated from flowering dates and previous observations on the same genotypes. After four weeks of treatment (at T1), clusters were sampled from the second set of plants entered into the PhenoDyn phenotyping platform. Once harvested, clusters of T0 and T1 samples were immediately photographed on both sides and weighed, before weighing 25-35 berries and individually wrapping them in aluminium foils for subsequent freezing in liquid nitrogen. Individually wrapped berries and the remaining part of the cluster were then stored at −80°C and −20°C, respectively.

One cluster per plant was photographed every 2 days during the whole treatment period, as in Savoi *et al*. (2021). Pictures of the 9 red genotypes were analyzed for projected areas of visible berries and for the percentage of green vs. non green pixels, with the Image J software (1.53t, java1.8.0_172, https://imagej.net/software/fiji/downloads). For this, the cluster was isolated on a white background (adjust color threshold) before converting to HSB stack and analysing hue angle histogram. The boundaries of “green pixels” on the histogram were determined through separate analysis of green berries.

#### 1.3. Organic acid and sugar analyses

Organic acids, glucose and fructose were extracted and measured by HPLC, as in Bigard *et al*. (2022), from cluster juice for all genotypes, as well as from individual berries for six selected genotypes.

#### 1.4. Statistical analyses

Data were analyzed with EXCEL® and R Software version 3.3.5 (R Core Team, 2021). Analysis of variance was performed on the surveyed variables, followed by the Tukey test (p<0.05). A loess curve was fitted (ggplot package, geom_smooth() function with span between 0.7 and 0.75 as indicated in figure legends) to obtain the trend of variables against irrigation level or time.

## Results and Discussion

### 1. Effects of water stress on the average sugar and malate contents at the usual fruit population level

During the 4-week experiment, sugar concentration showed an overall increasing trend from T0 to T1, accompanied by a decrease in malic acid (Figure 1), which confirms that ripening could actually be induced in all genotypes after the expected véraison date, as confirmed by colour analysis (Figure S4). For half of the genotypes, glucose plus fructose concentration even reached 1 M in at least one treatment, indicative of a clear ripe stage in less than four weeks (Bigard *et al*., 2018). However, high water availability (a to d treatments) unexpectedly led to a nearly complete inhibition of ripening for ‘VDQAG14’, ‘23298Mtp7’, ‘Arbane’, ‘7G31’ and ‘Opsimo Edessis’. To the best of our knowledge, complete inhibition of ripening in irrigated conditions has never been reported outdoors – though López *et al*. (2009) suggested a delay in the induction of ripening in these conditions. Typical features of the ripening process were observed for other genotypes and conditions, namely softening (not shown), sugar accumulation and subsequent growth resumption, malate breakdown and, if applicable, reddening (Figures 1, 2, 3, S4). However, there was large variation in berry relative expansion between T0 and T1, depending on the genotype (Figure 2). Berry weight could more than double for ‘VDQAG5’, ‘Savagnin blanc’ and ‘Arbane’, while growth was much more limited for ‘Cococciola’, ‘Garrido Macho’ and ‘7G39’. On average for all cultivars in the absence of strong WD (a to g treatments), berry expansion at T1 did not really start before reaching a minimal 0.4 M sugar concentration threshold, and was quite proportional to sugar concentration until reaching 1 M sugar, after which shrivelling occured (Figure S5). Severe WD conditions strongly inhibited berry growth in most genotypes. Nonetheless, growth data were scattered, as expected from the intrinsic variability of average berry weight from one cluster to the other (Shahood *et al*., 2020). This variability inevitably translated into noise on sugar accumulation, calculated as the product of sugar concentration x volume (itself approximated by berry weight) (Figure S1).

**Figure 1.**
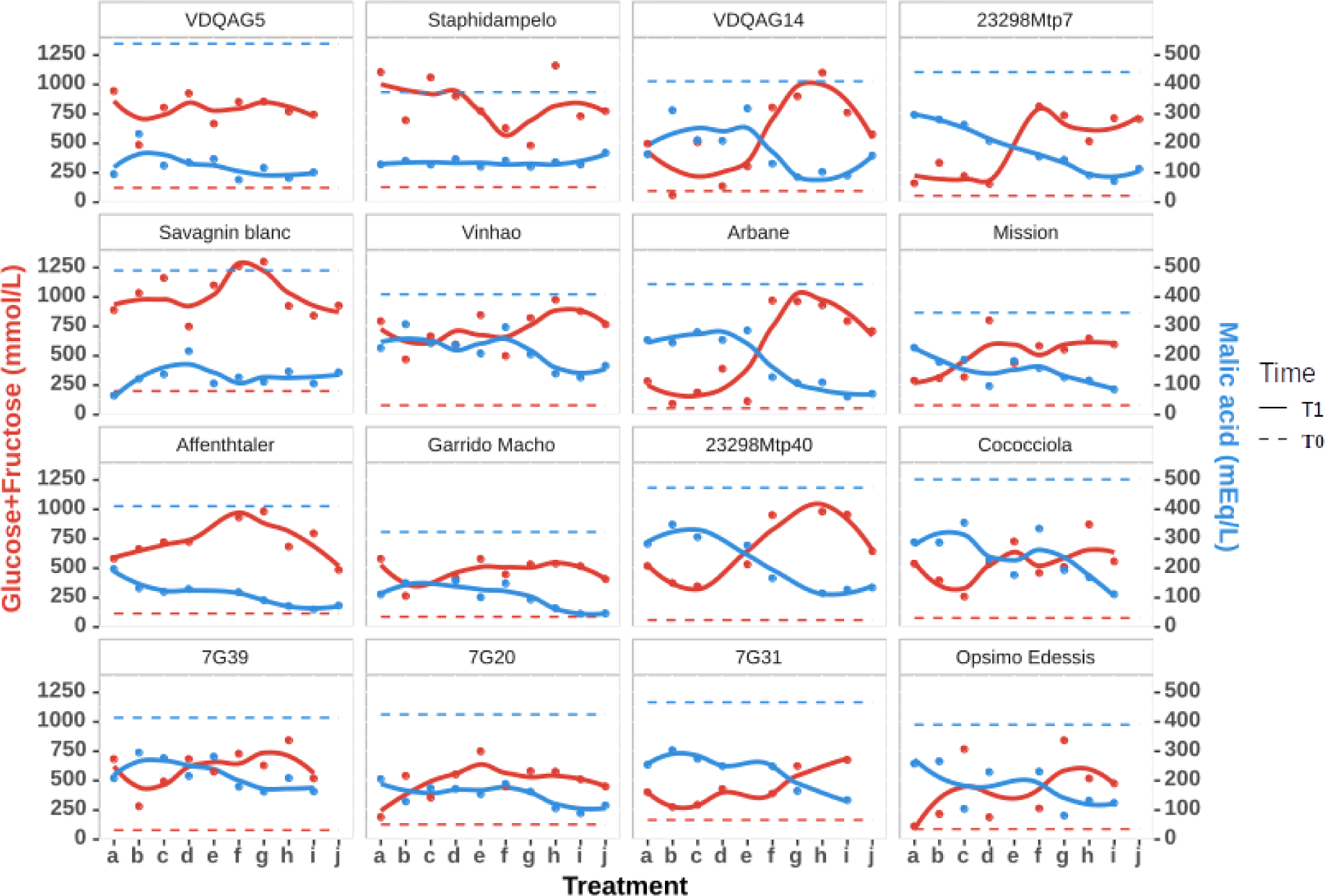
Effect of water deficit on the concentration of sugar and malic acid as classically measured on samples of averaged berries. At T0, 5 plants were available, allowing the composition of their respective clusters to be averaged (dotted lines). Only one plant per treatment was available at T1. T0: a few days (3-4 days) before the expected véraison stage; T1: 28 days after T0. Treatments: a, WW; b, c, d, slight WD; e, f, g, medium WD; h, i, j severe WD. The continuous line is a loess fitting (span = 0.75).

**Figure 2.**
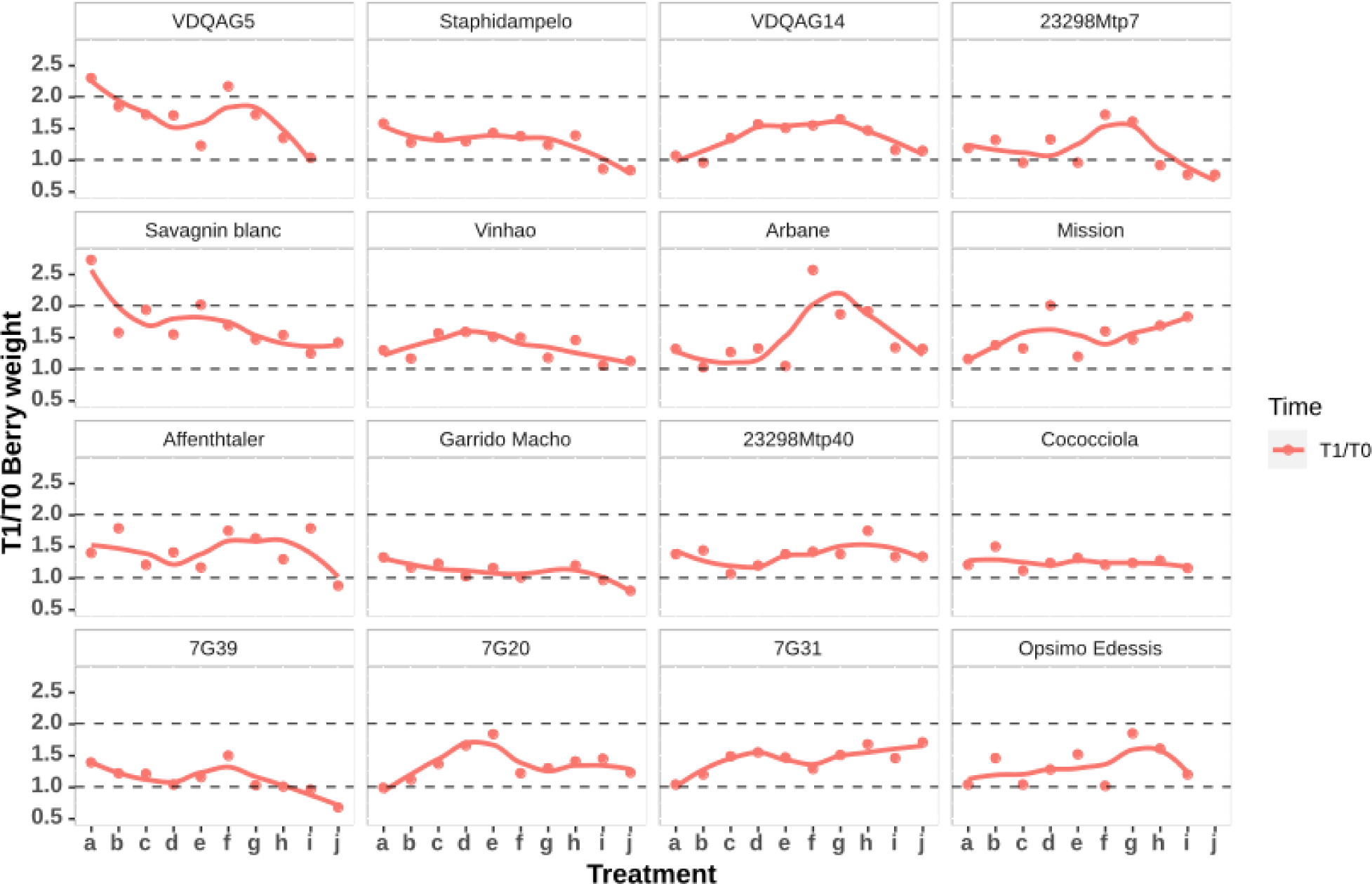
Effect of water deficit on berry expansion, as classically measured at average berry population level. 25-35 berries per plant were weighed to calculate the T1/T0 expansion ratio (averaged over 5 plants per genotype at T0, a few days before expected véraison date; from one plant per genotype at T1, 28 days after T0). Treatments: a, WW; b, c, d, slight WD; e, f, g, medium WD; h, i, j severe WD. The continuous line is a loess fitting (span = 0.70).

To overcome this issue, we normalized the data using tartrate concentration. As the quantity of tartrate per fruit remains fairly constant after véraison (Burbidge *et al*., 2021; Rienth *et al*., 2016; Rösti *et al*., 2018; Ruffner, 1982; Terrier *et al*., 2001), the concentration ratio of any solute to tartrate provides a valuable proxy for the quantity of solute per sample (see rationale in Supplementary material). It shows that on average and regardless of the capacity to ripen from a to d treatments, sugar accumulation was often maximal for the f or g treatments and dramatically impaired at the strongest WD levels. Malate breakdown was extremely limited in the five genotypes that did not accumulate sugars from a to d treatments (compare Figure 3 to Figure 1). As already discussed in Bigard *et al*. (2018, 2022) in the WW case, malate (Figure 1), tartrate (Figure S2) and the malate/tartrate ratio (Figure 3) displayed genotypic differences at the end of the green stage (T0), with ‘VDQAG5’ showing the highest malic acid level and ‘Garrido Macho’ the lowest one. We not only confirm here that WD promoted malic acid breakdown, in accordance with Zheng *et al*. (2018) and López *et al*. (2007), but also show that WD uncoupled it from sugar accumulation (Figure 3).

**Figure 3.**
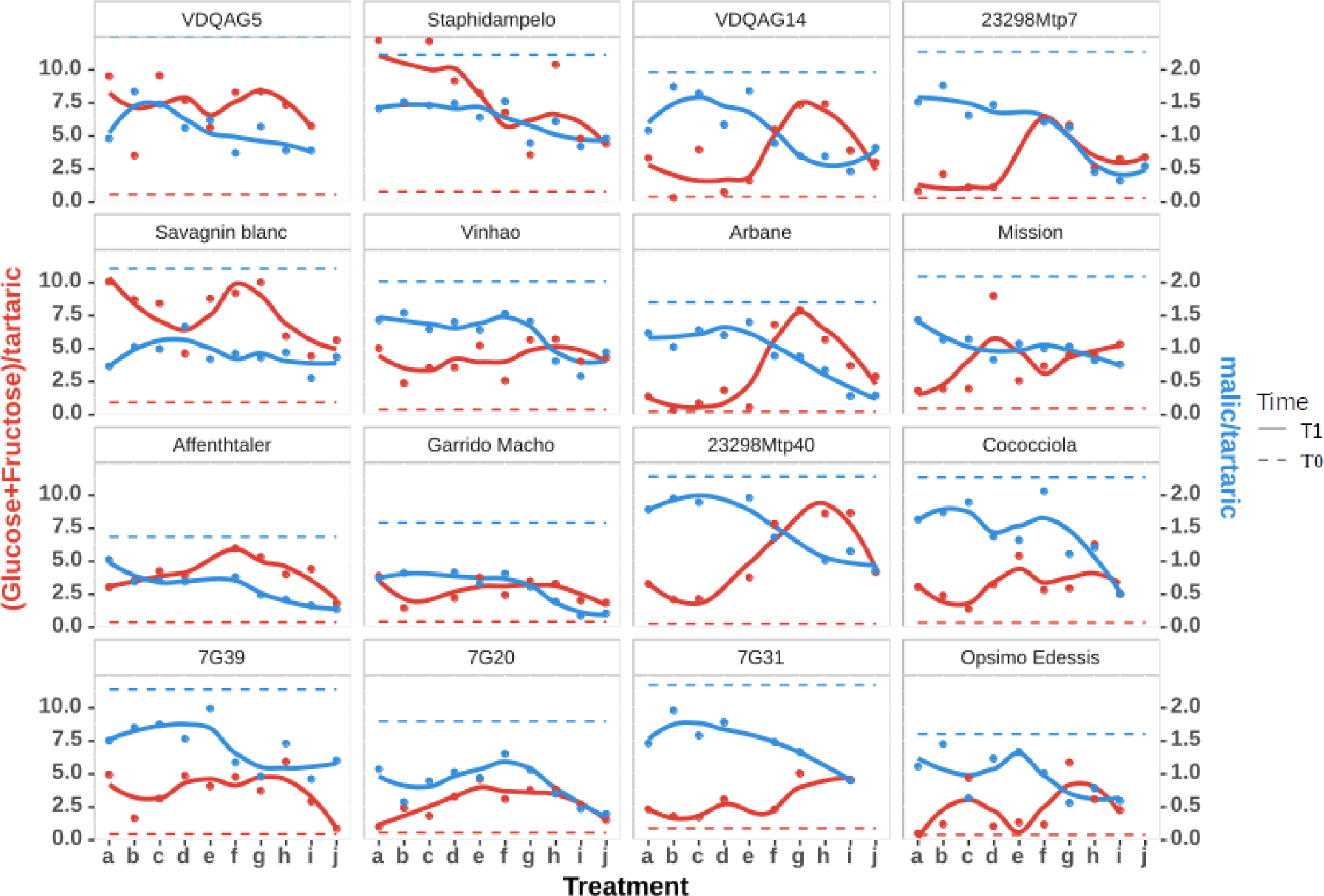
Effect of water deficit on sugar accumulation and malic acid breakdown as assessed with the tartrate ratio at average berry population level. At T0, 5 plants were available, allowing the composition of their respective clusters to be averaged. At T1, only one plant per treatment was available. T0: a few days (3-4 days) before the expected véraison (softening) stage; T1: 28 days after T0. Treatments: a, WW; b, c, d, slight WD; e, f, g, medium WD; h, i, j severe WD., The continuous line is a loess fitting (span = 0.75).

### 2. Effects of water stress on single berry ripening

Six genotypes approaching more than 0.8 M sugars four weeks after the anticipated veraison date were selected for single berry analysis on 25 to 35 berries per treatment (‘VDQAG5’, ‘23298Mtp40’, ‘Affenthaler’, ‘23298Mtp7’, ‘Arbane’, ‘VDQAG14’). The inhibition of sugar accumulation and growth by severe WD was striking (Figure 4), despite the aforementioned variability in average berry weight per cluster that is largely determined from the green stage onwards (Shahood *et al*., 2020). In WW condition, the variability in individual sugar concentration (obviously, the angle of each point in polar coordinates in Figure 4) was generally higher than that of berry weight, except when ripening was totally inhibited (‘23298Mtp7’) or nearly achieved (‘VDQAG5’). In this latter situation, berries from a and f treatments clustered together, close to the 1 M diagonal that is quite systematically reached with medium WD (f treatment). Although the berries with the lowest sugar concentration were also the smallest ones, as can be expected for late-ripening berries still expanding while accumulating sugars. However, a treatment excepted, intra treatment variations in the sugar content were largely dominated by huge erratic variations in berry volume at quite stable concentration (Figure 4). The strongest WD (treatment j) resulted in limited sugar concentration and berry volume, suggesting a considerable inhibition or delay of phloem unloading into each individual berry, thus excluding the simultaneous presence and competition between “normal ripening berries” and “water and sugar limited ones”.

**Figure 4.**
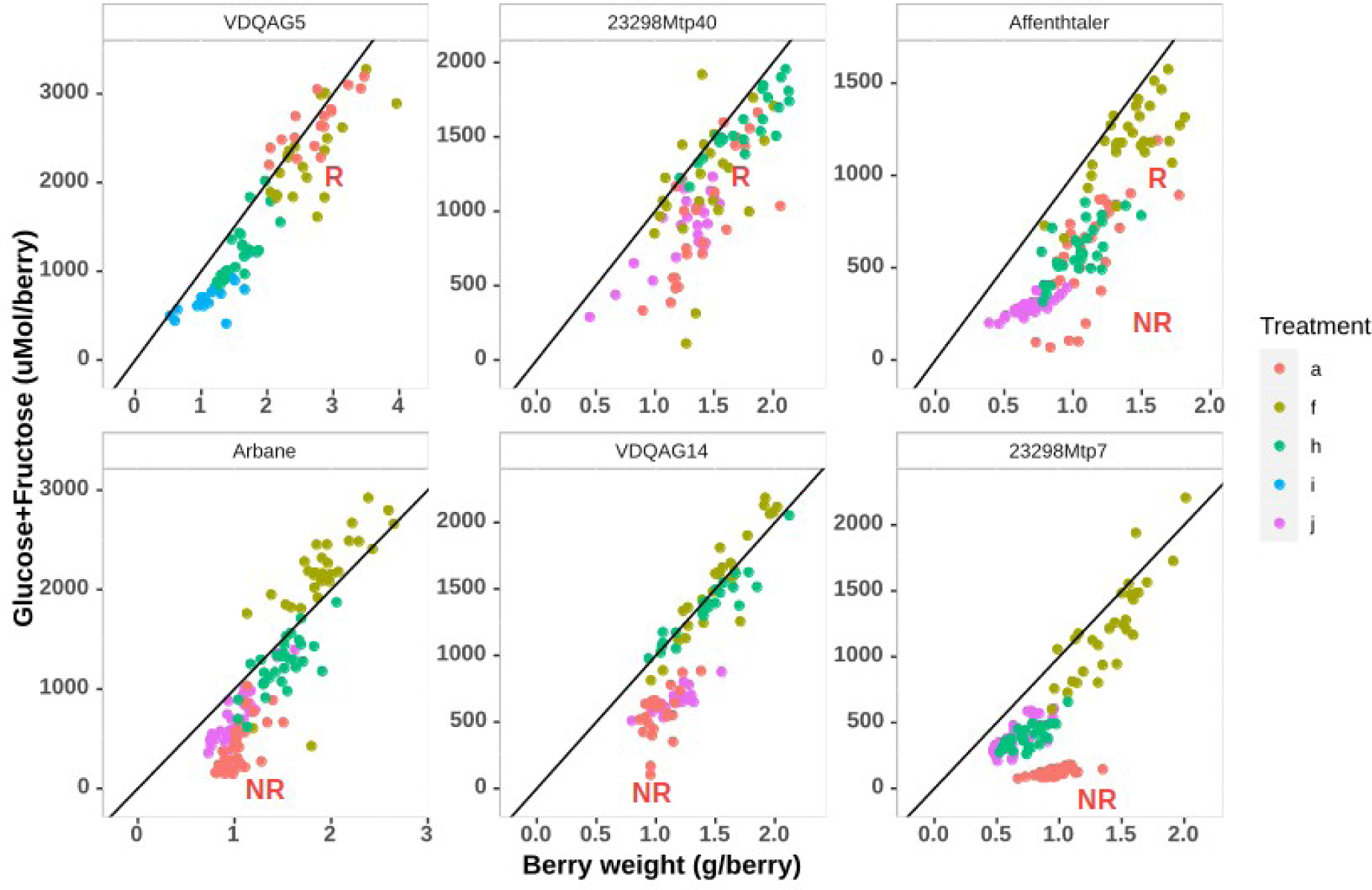
Effect of water deficit on individual berry weight and the quantity of sugars accumulated in single berries at T1. The black line refers to 1 M sugar concentration (assuming berry density is 1). Treatments: a, WW; f, medium WD; h, i, j severe WD. R: single berries exhibiting usual ripening in fully irrigated conditions (treatment a). NR: single berries unable to ripen in fully irrigated conditions. For some of these genotypes part of the berries started to ripen in fully irrigated conditions (‘23298Mtp40’, ‘Affenthaler’, ‘Arbane’, ‘VDQAG14’).

As a general rule, berry weight was strongly impaired between the f and j treatments, suggesting a complete collapse of the second growth period in the most stringent conditions, except for ‘23298Mtp40’ (Figure 4). However, j-treated berries exhibited all signs of ripening, namely softening, colouring, and accumulating sugar. Using tartaric acid concentration as a proxy for berry expansion (see above) evidenced that in the most stressed i and j conditions, more or less intense shrivelling of individual berries took place during the 28 days treatment (Figure S3), possibly masking complex growth dynamics.

### 3. Effects of water stress on ripening kinetics

We tried to quantify the growth and coloration kinetics simultaneously through time-lapse photography. For ‘23298Mtp7’, cluster surface increased with time under g treatment, and diminished under j treatment (Figure 5), consistent with T0/T1 metabolic data for this genotype (Figures 2, S1, S3). More than two weeks were needed to trigger complete cluster coloration under g treatment, which was achieved in only four days under severe stress, highlighting a better synchronisation of berries in severe WD conditions, which could be confirmed in all other cultivars except Vinhao (Figure S4). Figures 5 and S4 also show that a variable delay occurred between T0 and the onset of ripening among genotypes and treatments.

**Figure 5.**
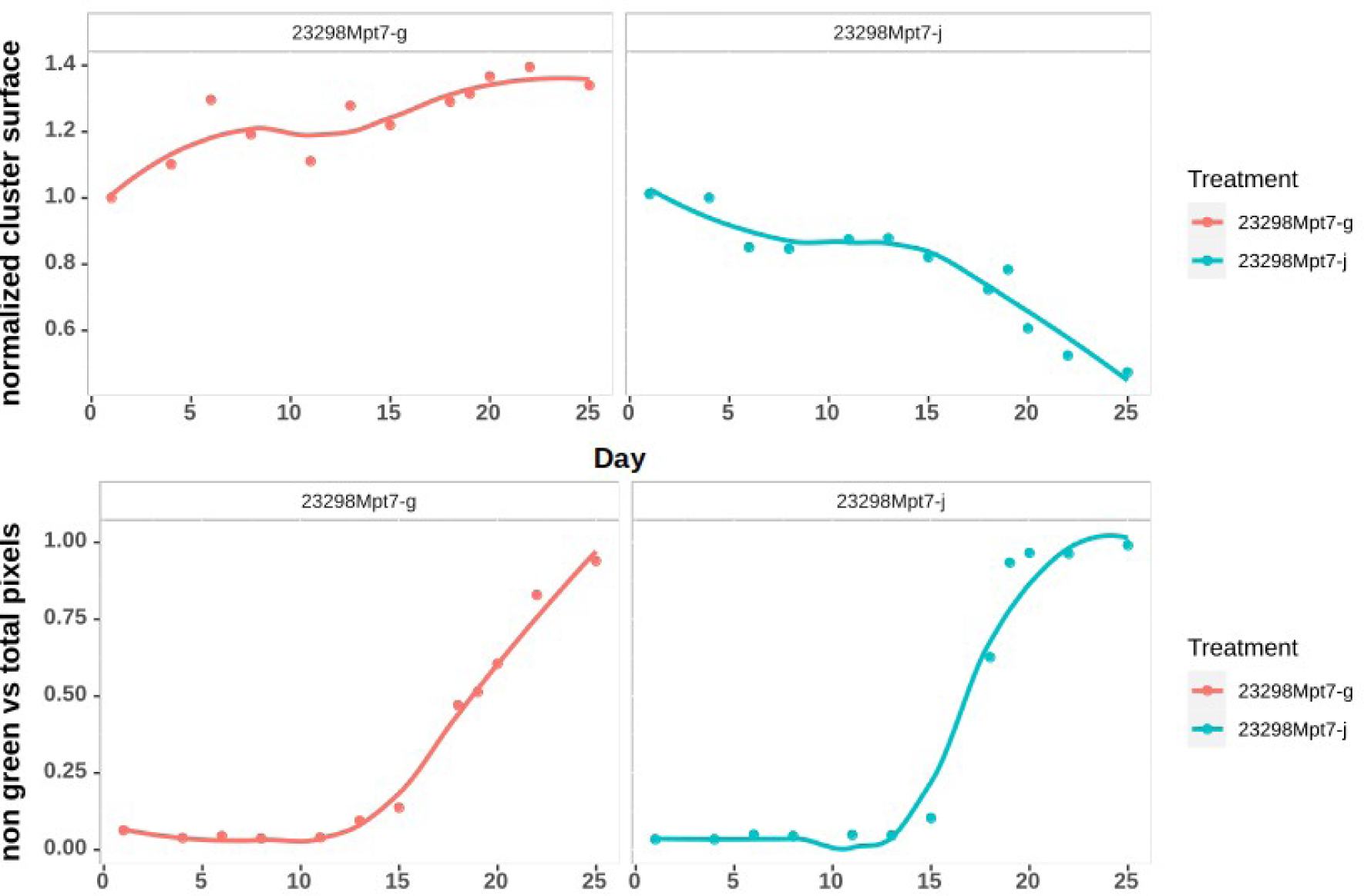
Dynamics of cluster growth and coloration from T0 to T1, as appreciated from image analysis. Cluster surface refers to the normalized (divided by T0 pixels) count of non-background pixels and color index to the proportion of non-green pixels. The continuous line is a loess fitting (span = 0.8) Treatments: g, medium WD; j severe WD.

Therefore, we also attempted to distinguish between phenologic and metabolic effects of water deficit, taking advantage of the natural asynchronicity of single berries. As presented above, changes in the dilution of tartaric acid are a convenient proxy to identify the respective phases of berry expansion or shrivelling. We know that sugar unloading in berries stops at their maximal berry volume or minimal tartaric acid concentration, after which variations in water content (e.g. water loss during shrivelling) may affect the concentrations of both solutes simultaneously (Bigard *et al*., 2022; Savoi *et al*., 2021). This results in a ‘dilution diagonal’ (passing through the origin) typical of the shrivelling stage as illustrated in Figure 6. This figure takes advantage of our previous exhaustive characterization of ‘VDQAG14’ single berries, in 2016 and 2017 in a well-watered vineyard (Bigard *et al*., 2020), to benchmark present greenhouse experiment. In our present experiment, ‘VDQAG14’ exhibited an overall ‘delayed ripening’ pattern in well-watered condition with the ripest fruits in a highly asynchronous population only reaching 0.7 M hexose (Figure 6B) and partial expansion. Individual berries from the a and j treatments grouped at low sugar and high tartaric acid concentrations, as opposed to other treatments. However, the distribution of WW berries indicates that they were still in their expansion phase, with tartaric acid dilution increasing with sugar concentration. In contrast, j-treated berries were distributed along a ‘dilution diagonal’, as if extra sugar concentration was principally a matter of impaired expansion and water accumulation, rather than accelerated sugar accumulation. Berries from medium WD treatments (f, h) actually reached sugar concentration around 1 M, as expected following a 28-day ripening period (Shahood *et al*., 2020). Even though less diluted than outdoors, these berries also exhibited a proportional increase in sugar and tartrate concentrations, indicating they certainly just reached the post-phloem arrest shrivelling period.

**Figure 6.**
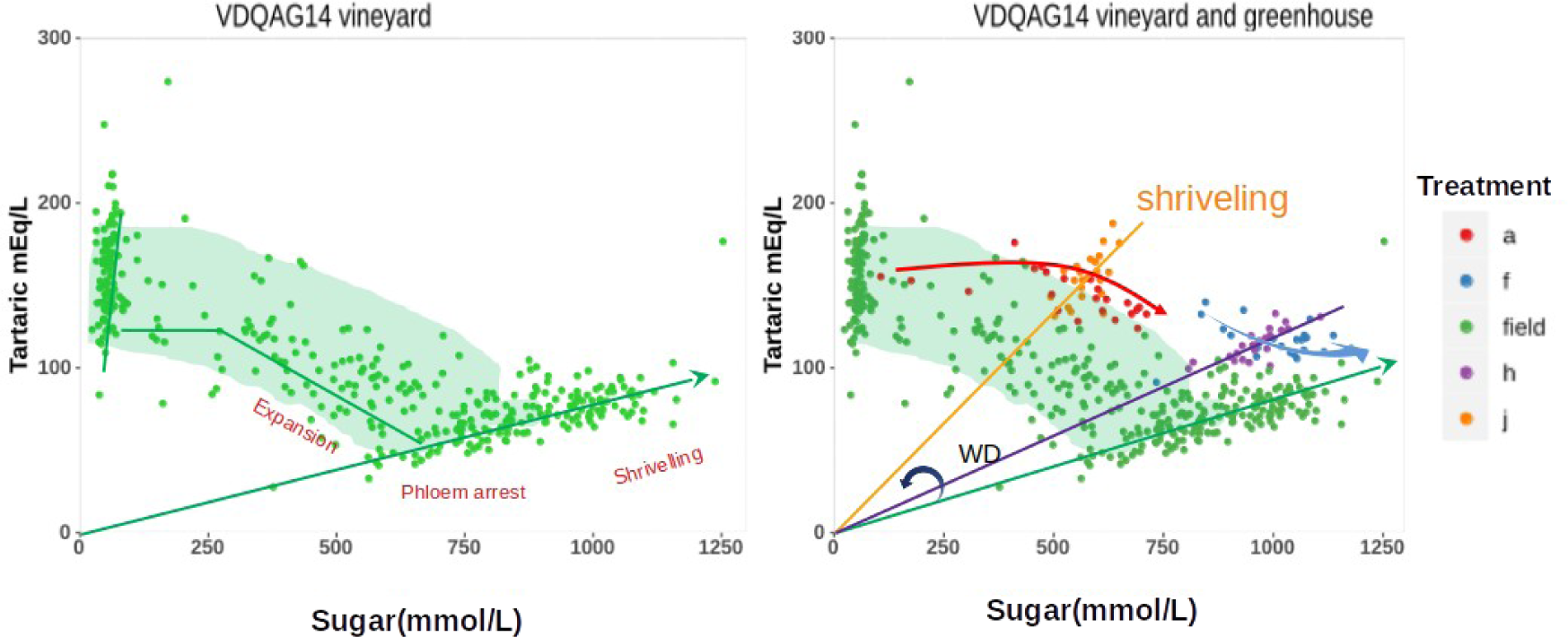
Single berry expansion, sugar concentration, and phenology according to water deficit. (A) Berry developmental phases according to tartrate dilution (1 point = 1 berry, sampled from flowering to over-ripening) in a standard vineyard (data from Bigard *et al*., 2021). (B) Same as A with in addition berries sampled 28 days after the anticipated date of véraison, following full-watering (treatment a), or medium (f) to severe (h, j) water deficit. Berries on each straight line passing through the origin may only differ by dilution unless other metabolites indicate the opposite. The advancement of ripening can be assessed by the clockwise angle with the vertical axis.

## General discussion

As cell division definitively stops largely before véraison (Ojeda *et al*., 2015), the inhibition of the second growth period reported here under severe WD necessarily resulted from reduced cellular expansion during ripening.

As in other non-climacteric fruits, grapevine berry ripening is a highly regulated developmental process that requires switching on a diverse set of genes encoding membrane transporters and cell wall hydrolases, recruited for activating both the apoplastic pathway of sucrose unloading from phloem and vacuolar expansion (Savoi *et al*., 2021). Water transport by aquaporins being passive, a descending overall water potential gradient is also a strict prerequisite for the expansion of the pericarp, which is achieved through the accumulation of major osmolytes. It is empirically observed that berry volume progressively doubles as hexose concentration reaches 1.0 M in usual conditions, as globally confirmed here on genetically distant genotypes. Berry expansion first depends on its capacity to induce softening and sugar accumulation, which was inhibited under WW in some cultivars (Figures 2, 3, 4). However, abrupt softening and relaxation in turgor pressure marking the dramatic acceleration of sugar loading at the onset of ripening are not enough to trigger the second growth period, which starts only when reaching ca 0.4 M hexose or 0.55 osmol (Shahood *et al*., 2020, supplementary figure S5). Such an osmotic threshold could be indicative of a 0.5 M sucrose concentration in the phloem, precisely the one at which calculated xylem backflow vanishes in growing berries (Keller & Shrestha., 2014). We show here that the induction of ripening and sugar accumulation is resilient to some level of WD. Moderate WD could even be qualified as favourable to ripening, as well-watered conditions in the greenhouse hindered the triggering or synchronisation of ripening in several genotypes. This is in line with the recent suggestion that ripening speed increases when Ψb decreases up to −0.8 MPa, and slows down at more severe Ψb (van Leeuwen *et al*., 2023). It is tempting to speculate that such results reflect variations in the auxin/ABA balance, which respectively acts as inhibitor or promoter of ripening in grapevine (Böttcher *et al*., 2011; Davies & Robinson, 1996; Ferrer *et al*., 2016). Additionally, with strongly reduced root volume, limited evaporative demand (relatively low irradiance and VPD) and attenuated daily amplitudes of temperature, greenhouse conditions noticeably differ from vineyard ones, in which full irrigation does not block ripening. Reduced cluster transpiration can slow down ripening (Keller *et al*., 2015), so limited evaporative demand in the greenhouse might also explain why some genotypes showed inhibited ripening in WW conditions.

Under intermediate WD, all genotypes exhibited near-optimal berry growth and sugar accumulation, whereas increasing WD progressively impaired both processes, in line with the results of de Almeida *et al*. (2023). The 10% rise in sugar concentration reported by Bécart *et al*. (2022) following a fifty years study in the Rhone Valley parallels the impairment of berry volume at technological maturity, thus the process of sugar accumulation into the fruit was kept rather unaffected by climate change in this region up to now. Such increase in sugar concentration due to greater impairment of final berry volume than of sugars accumulation was not detected in our greenhouse conditions. When compared to outdoors conditions, in addition to previously cited reasons, clay was less abundant in the substrate (10 % volume), forcing soil water potential to oscillate in phase with irrigation, rather than to progressively reach increasing negative values with decreasing water availability. Berries were also deliberately harvested only 4 weeks after véraison at best, in order to focus on the phloem unloading period and limit as far as possible the confusing effects related to the post-phloem arrest shrivelling period, which, since harvesting 2 weeks later, is an integral part of the technological ripening process. Whatever, field studies have also shown a dramatic reduction in berry expansion and sugar accumulation under intermediary to severe water stress, as evidenced here (Gambetta *et al*., 2020; Yang *et al*., 2022). In contrast, Hewitt *et al*., (2023) just reported a comparable experiment in own rooted vines in climatic cabinets, in which berry growth, malic acid and sugars were kept unaffected by a later (one week post softening) and shorter WD (one week, 50% field capacity, midday leaf water potential −1.3 MPa). Clearly however, as berries reached 18-20 brix (ca 1 M sugars) after one week treatment, the osmotic pressure was favourable to water accumulation, and berries already expanded at maximal rate, when challenged by the stress.

The intensity of malate breakdown proved more sensitive to WD than that of sugar accumulation. Shahood *et al*. (2020) and Bigard *et al*. (2022) put forward that, under non-stressful conditions, roughly 4 hexose molecules should be accumulated per disappearing malic acid during the first phrase of ripening, in line with the induction of the probable sucrose/H^+^ antiporter *VvHT6* (Savoi *et al*., 2021). Malic acid is progressively exhausted and replaced by sugars as a respiratory substrate later on the sugar accumulation process, while H^+^ pumps progressively replace malate efflux in the electro-neutralisation of the sugar/H^+^ antiport. Savoi *et al*., 2021 noticed that the overall malate breakdown/sugar accumulation ratio was impaired, indicating greater ATP requirements, and that there was increased transcription of the energy requiring plasma membrane sugar H^+^ symporter *VvHT2* in external fluctuating conditions. Patono *et al*. (2022) evidenced that, in line with sugar influx, the expression of *VvSWEET10* and *VvHT6* genes, both key players for berry sink strength, is repressed upon WD, and reactivated following rewatering. Thus, a picture emerges suggesting that energy consuming transport processes are activated upon stress, to counterbalance a restricted sugar supply. Further transcriptomic studies are needed for a better understanding of the molecular bases of the acceleration of malate breakdown upon WD.

## Conclusion

Severe water limitation led to huge inhibition of the second berry growth period and to variable levels of shrivelling among individual fruits, but not in the onset of sugar accumulation and the synchronisation of coloration, while véraison was delayed or even prevented in some genotypes under well-watered conditions. Although the quantity of sugar per fruit was drastically reduced, the sink strength for photoassimilates remained considerable even at severe WD conditions entailing partial leaf shedding, fruit growth inhibition, and berry shrivelling. Such results question phloem mass flow and sugar concentration under severe WD.

Limited WD was quite systematically beneficial to the ripening process, in terms of individual berry growth and sugar accumulation, or quite neutral for those cultivars that were able to ripen in well-watered conditions. From a methodological point of view, while highlighting difficulties in modelling vineyards environmental conditions under glass, our study emphasises the need for real-time single berry growth monitoring (Daviet *et al*., 2023) to deconvolve the phenologic and metabolic consequences of environmental conditions on sugar and water fluxes.

## Acknowledgements

The authors thank the French national research agency (ANR<u>-19-CE20-0024</u>) and the Institut Agro Montpellier (Preciput ANR) for funding the G2WAS project. The PhD scholarship of MS was supported by ANR and the Occitanie region. The post-doc scholarship of SS was supported by ANR and the Poupelain Foundation. The authors also thank the DAAV (UMR AGAP Institut) and ETAP (UMR LEPSE) research groups for their participation in plant preparation, management and phenotyping.

